# Optimizing automated detection for cytoplasmic TDP25 aggregates in fluorescence imaging

**DOI:** 10.1101/2025.04.04.647339

**Authors:** Sumire Sogawa, Kotetsu Sasaki, Akira Kitamura

## Abstract

Protein aggregates are known to disrupt normal cellular functions and homeostasis, serving as key hallmarks of various neurodegenerative disorders, including Alzheimer’s disease, Parkinson’s disease, and amyotrophic lateral sclerosis (ALS). Automated detection of cytoplasmic, disease-associated aggregates in fluorescence images is crucial for characterizing these aggregates and exploring potential strategies for their prevention. In this study, we demonstrate that removing background fluorescence and improving the brightness of aggregates significantly enhances the detection efficiency of cytoplasmic aggregates formed by the 25 kDa C-terminal fragment of ALS-associated TDP-43 (TDP25) using an automated aggregate detection algorithm. A high signal-to-noise ratio can improve detection efficiency. Our findings contribute to the development of more effective detection methods for disease-associated aggregates of heterogeneous sizes and fluorescence intensities, which are typically challenging to identify automatically.

## Introduction

Misfolded and unfolded proteins accumulate with other proteins and biomolecules to form oligomers, multimers, and large aggregates [1,2]. Genomic mutations, cellular stress, and imbalance in protein quality control systems—including nascent polypeptide synthesis, folding, and degradation— promote to the formation of aggregate from misfolded/unfolded proteins. Protein oligomers and multimers are not always localized as puncta; they can also diffuse [3,4]. In contrast, because aggregates are often biochemically insoluble, they can appear as bright spots (puncta) within cells [3]. Other intracellular punctate structures include condensates, which form via liquid-liquid or liquid-solid phase separation owing to weak, yet multivalent interactions, reaching an equilibrium state with two different phases [5,6].

In various neurodegenerative disorders, such as Alzheimer’s disease, Parkinson’s disease, and amyotrophic lateral sclerosis (ALS), condensates and aggregates have been observed in neurons derived from patients. These aggregates of disease-associated proteins can disrupt normal cellular functions and homeostasis, resulting in cytotoxicity and contributing to the progression of neurodegenerative disorders from a neurotoxicity perspective [7]. Understanding the characteristics of protein aggregates is crucial in cell biology, medicinal chemistry, and biotechnology, particularly for their prevention and clearance. For example, when investigating anti-aggregation drugs, a key step is evaluating whether the proportion of intracellular aggregates decreases following drug administration by recognizing intracellular puncta. Therefore, accurately counting the number of intracellular puncta with minimal effort is important.

Many studies have reported automated puncta detection algorithms using fluorescent images [8-13]. One recently developed algorithm, PunctaFinder, not only detects puncta but also defines cytoplasmic regions [14]. PunctaFinder automatically provides a more accurate number of bright spots compared to other puncta-detection algorithms, such as TrackMate, μ-Track, and SpotiFlow [14]. We have previously reported on the formation mechanism and toxicity of ALS-associated cytoplasmic aggregates of a 25 kDa C-terminal fragment of TDP-43 (TDP25; amino acids 220-414) [15-17]. Automated measurements are challenging due to the wide variability in the shape, size, and number of TDP25 cytoplasmic aggregates across different cells. However, PunctaFinder, which detects cytoplasmic regions, may offer an advantage in automatically detecting this variety of TDP25 cytoplasmic aggregates.

In this study, we verified that PunctaFinder is an effective algorithm for automatically detecting cytoplasmic TDP25 aggregates. While PunctaFinder alone did not achieve a particularly high detection efficiency for TDP25 aggregates in immortalized adherent cells, the detection accuracy improved when background fluorescence intensity was appropriately removed and brightness was amplified. Furthermore, it was demonstrated that the same approach allowed efficient detection of TDP25 aggregates in nematode embryos. The results of this study make a significant contribution to the development of a highly efficient detection procedure for disease-associated aggregates with heterogeneous sizes and fluorescence intensities, which are typically difficult to automate.

## Materials and Methods

### Cell culture, transfection, and image acquisition

Murine neuroblastoma Neuro-2a cells (#CCL-131; ATCC, Manassas, VA, USA) were cultured in DMEM (D5796; Sigma-Aldrich, St. Louis, MO, USA) supplemented with 10% FBS (12676029, Thermo Fisher Scientific, Waltham, MA, USA), 100 U/mL penicillin G (Sigma-Aldrich), and 0.1 mg/ml streptomycin (Sigma-Aldrich), as previously described [17]. Plasmid DNA for green fluorescent protein (GFP)-tagged TDP25 (G25), with GFP tagged at the N-terminus, was used as previously described [15,16]. Cells (2.0 × 10_5_) were plated in a 35 mm glass-bottom dish (3910-035; IWAKI-AGC Technoglass, Shizuoka, Japan) 1 day before transfection. Plasmid DNA (1 μg for G25 expression) was transfected using Lipofectamine 2000 (Thermo Fisher). After overnight incubation for transfection, the medium was replaced with fresh medium, and the cells were examined under microscopic observation. Cells were observed using an LSM 510 META (Carl Zeiss, Jena, Germany) with a Plan-Apochromat 10×/0.45NA M27 objective (Carl Zeiss) for puncta detection, or a C-Apochromat 40×/1.2NA Korr. UV-VIS-IR water immersion objective for typical images shown in Figure 1a, all maintained at 37°C. GFP was excited using a 488 nm wavelength incident laser, and the excitation beams were divided using a HFT405/488. Fluorescence was detected with a photomultiplier tube after passing through a 505 nm long-pass filter (LP505). Bright-field images were captured using a photomultiplier tube for transmitted light, attached to an inverted microscope Axioobserver.Z1 (Carl Zeiss). The pinhole size was set to its maximum. The image resolution was 2048 × 2048 pixels (X × Y), with a scanning speed of 1.6 μs per pixel. The acquired images of cells with G25 aggregates were then used for subsequent image analysis.

**Figure 1.**
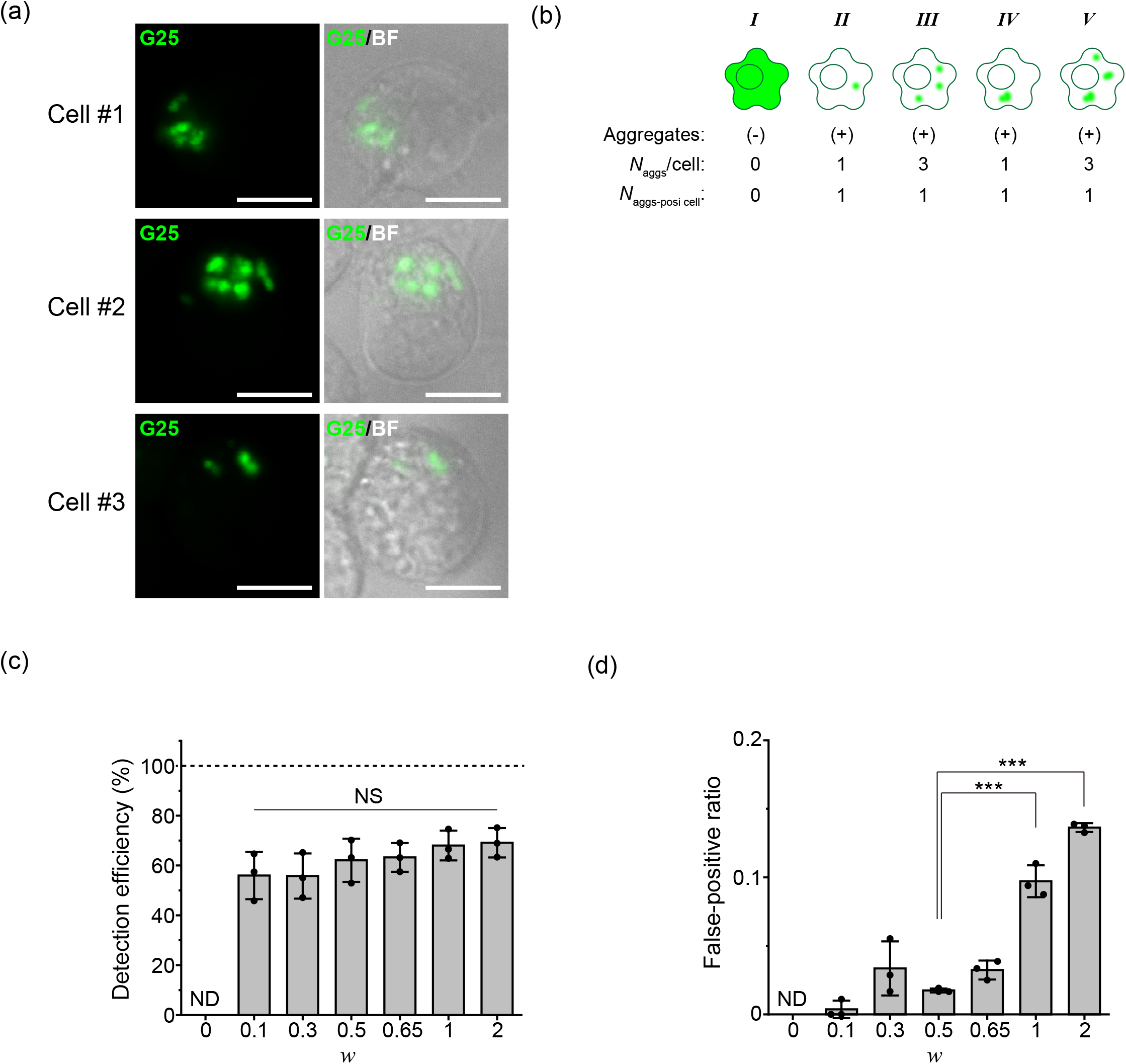
Issues in automatic detection of Neuro2a cells harboring TDP25 aggregates in the cytoplasm. (a) Three fluorescence and bright-field images of typical Neuro2a cells harboring aggregates of GFP-TDP25 (G25) in the cytoplasm as multiple punctate structures. The images show GFP fluorescence alone (left) and an overlay of bright-field (BF) and GFP fluorescence (right). Bar = 10 μm. (b) Example illustration of cells with and without aggregates ((+) and (-), respectively). In cells without aggregates, GFP-TDP25 is uniformly distributed throughout the cells (*I*). In contrast, GFP-TDP25 aggregates appear as highly fluorescent, punctate structures in the cytoplasm (*II*-*V*). The number of aggregates per cell (*N*_aggs_/cell) and the number of aggregate-positive cells (*N*_aggs-posi cell_) corresponding to each of the cells in the top illustration are shown. Aggregates that could not be distinguished separately in the fluorescence images because of the limitation of optical resolution were counted as a single aggregate (*IV* and *V*). (c) Detection efficiency of cells harboring aggregates of GFP-TDP25 in the cytoplasm with varying weighting factor (*w*). The bars and dots show the results from three independent trials (mean ± SD; n = 3). ND: No aggregates were detected automatically using the PunctaFinder algorithm. NS: Not significant (*p* ≥ 0.05). (d) The ratio of false-positive puncta automatically counted to the number of aggregates manually counted when changing the weighting factor (*w*). The bars and dots show the results from three independent trials (mean ± SD; n = 3). ND: No aggregates were detected automatically using the PunctaFinder algorithm. Statistical significance: \*\*\**p* < 0.001.

### Nematode culture and image acquisition

Nematodes (*Caenorhabditis elegans*) were maintained and handled according to standard methods, cultured at 20°C [18]. Transgenic animals expressing G25 in the body wall muscles under the regulation of the *myo-3* promoter and enhancer elements from the *unc-54* myosin heavy-chain locus were of the same strain as describd in a previous report [17]. The animals were grown at 20°C on nematode growth medium (NGM) agar plates seeded with *Escherichia coli* OP50 strain (Caenorhabditis Genetics Center, Minneapolis, MN, USA) as a food source. Fertilized embryos were recovered from adult animals in M9 buffer and dissected under a dissection microscope (Stemi 305; Carl Zeiss, Jena, Germany). Embryos in M9 buffer were transferred to the wells of a glass-bottom dish (3970-103, IWAKI, Shizuoka, Japan). Time-lapse observation of embryogenesis and aggregate formation were performed using an LSM 510 META with a C-Apochromat 40×/1.2NA W Korr UV-VIS-IR water immersion objective lens (Carl Zeiss) and immersion oil with the same refractive index as water (Immersol W 2010, Carl Zeiss), in a room maintained at 25°C. GFP was excited at a wavelength of 488 nm, and the excitation beams were divided using an HFT488/594. Fluorescence was detected using a photomultiplier tube after passing through a 505 nm long-pass filter (LP505). Bright-field images were captured using a photomultiplier tube for transmitted light, attached to an inverted microscope Axioobserver.Z1 (Carl Zeiss). The pinhole size was set to 71 μm (1.0 airy unit), and the image resolution was 2048 × 2048 pixels (X × Y) with a zoom factor of ×0.7. The scanning speed was 0.8 μs per pixel. Images were acquired every 5 min for 180 cycles. Embryo images with G25 aggregates were selected from the time-lapse series and analyzed.

### Image analysis

A custom-built computer running Microsoft Windows 11 was used for image analysis. The system consists of a PRIME B760M-A D4 motherboard (ASUSTeK Computer Inc., Beitou District, Taipei, Taiwan), a Core i5-13500 central processing unit (Intel, Santa Clara, CA, USA), two 16 gigabytes (GB) of double data rate 4 synchronous dynamic random-access memory (total 32 GB; Kingston Technology, Fountain Valley, CA, USA), 500 GB MVMe PCIe Gen4×4 solid state drive (ADATA Technology Co., Ltd., New Taipei City, Taiwan) for the operating system, and an 8,000 GB Serial ATA 3.0 7,200 rpm hard disk drive (Seagate Technology, Fremont, CA, USA) for data storage. Python (version 3.12.7) was executed using the Anaconda distribution (Conda version 24.11.1, Conda-build version 24.9.0; Anaconda Inc., Austin, TX, USA). The Jupyter Notebook environment for running PunctaFinder was obtained from the original developer’s GitHub repository (https://github.com/molecular-systems-biology/PunctaFinder; accessed April 1, 2025).

Following the procedure of the PunctaFinder algorithm [14], fluorescence images of G25-expressing cells were used to generate cell masks. The images were binarized using Fiji-ImageJ version 1.54p (https://fiji.sc/; accessed on April 1, 2025). The edges of the analyzed images were filled with a zero value using a 20-pixel-wide square frame, as indicated below, with a custom macro in Fiji-ImageJ.

~~~
width = getWidth();
height = getHeight();
margin = 20;
makeRectangle(margin, margin, width - 2 * margin, height - 2 * margin);
run(“Make Inverse”);
setForegroundColor(0, 0, 0);
run(“Fill”);
run(“Select None”);
~~~

Cell-segmented images were created for cells with areas exceeding 100 pixels using the “Analyze Particles” command, followed by two 2D binary filters (“Erode” and “Watershed” commands) in Fiji-ImageJ. Segments that were not successfully separated were manually divided using drawing tools. Finally, cell masks with independent intensity values for each cell segment were generated using a custom macro in Fiji-ImageJ, as described below.

~~~
run(“Set Measurements…”, “area mean standard min centroid center perimeter bounding
fit shape feret’s redirect=None decimal=3”);
run(“Analyze Particles…”, “size=0-Infinity show=Masks clear include”);
run(“Connected Components Labeling”, “connectivity=4 type=[16 bits]”);
~~~

The Puncta detection dataset for PunctaFinder was generated from fluorescence images of 20 cells positive for G25 aggregates. Using this dataset, the threshold values (*T*_local_, *T*_global_, and *T*_CV_) were validated by minimizing the test statistic (#TS_*i*_), as shown in Eq. 1.

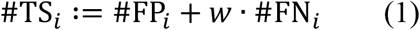

where #FP_*i*_ represents the number of false positives (FPs; verified non-punctate locations), #FN_*i*_ is the number of false negatives (FNs; verified punctate locations), and *w* is the weighting factor. Based on these criteria, G25 aggregates in Neuro2a cells and nematode embryos were automatically detected using PunctaFinder. Fluorescence intensity measurement, background subtraction, and digital intensity amplification were performed using Fiji-ImageJ software. Aggregate-positive cells were defined as those in which puncta were detected by Puncta Finder within the cell segments, after removing FPs. The ratio of the number of aggregate-positive cells automatically detected to the number manually counted was defined as the “detection efficiency.”

### Plots and statistics

Plots were generated using Microsoft Excel and Origin 2024 (OriginLab Corp., Northampton, MA, USA). One-way ANOVA followed by a post-hoc Tukey’s honest significant difference test was performed using Origin 2024 (OriginLab Corp.).

## 3. Results and Discussion

### 3.1. Identification of the weighting factor for TDP25 aggregates in PunctaFinder algorithm

In a stable human embryonic kidney cell line, which ensures homogeneous expression of G25 through a tetracycline-responsive repressor, G25 did not form aggregates in the cytoplasm [15]. Therefore, we transiently expressed G25 in Neuro2a cells to promote aggregate formation. As previously shown [15,16], G25 aggregates did not form as a single punctum but as multiple puncta in the cytoplasm (Figure 1a). Since these multiple G25 aggregates often overlapped due to the resolution limit of light microscopy, this study aimed to quantify the number of cells with aggregates rather than the number of puncta in the cytoplasm (Figure 1b). The detection efficiency was defined as the number of cells harboring G25 aggregates detected automatically using the PuntaFinder algorithm, divided by the number counted manually. In the original report of the PuntaFinder algorithm, it was recommended to keep the *w* less than 1 to minimize the detection of imaging noise as puncta. We examined whether cells harboring G25 aggregates could be detected automatically when *w* was varied from 0 to 2 in the microscope images (Figure 1c). When *w* = 0, cells with aggregates were not automatically detected (Figure 1c). As *w* was increased to 0.1, 0.3, 0.5, 0.65, 1.0, or 2.0, the detection efficiency did not change significantly and did not match the number of cells quantified manually (Figure 1c). The threshold values corresponding to these weighting factors are listed in Table A1. To determine whether the underestimation was due to a high contribution of FNs, *w* was increased to 2. However, the detection efficiency of aggregate-positive cells detected automatically did not change significantly compared to when *w* = 1.0 (Figure 1c). Furthermore, as a concerning observation, the proportion of FPs increased dramatically when *w* was set greater than 1 (Figure 1d). Therefore, it was confirmed that *w* < 1 is more effective at reducing FP. Although the lowest proportion of FPs was observed when *w* = 0.5 in the images used here (Figure 1d), the detection efficiency of aggregate-positive cells detected automatically at *w* = 0.5 was remarkably lower than that quantified manually (Figure 1c). In summary, since many aggregates were undetected, it is important to identify conditions that can make these undetected aggregates detectable. For the following analysis, a weighting factor of 0.5 was used.

### 3.2. Increase in the detection efficiency of the aggregate-positive cells by improving the signal-to-noise ratio of the fluorescence images

Since PunctaFinder is an algorithm that determines puncta based on fluorescence intensity classification, we compared the fluorescence intensities of detected and undetected G25 aggregates. The automatically undetected G25 aggregates showed a lower fluorescence intensity distribution compared to the detected G25 aggregates (Figure 2a). Increasing the apparent fluorescence intensity by multiplying it did not improve the detection efficiency (Figure 2b). This may be due to the similarity in fluorescence intensity between areas without fluorescence-positive cells (background) and aggregates with low fluorescence intensities. Next, the fluorescence intensity was amplified by subtracting the background fluorescence intensity. We compared the constant background subtraction procedure (CBS) and the rolling ball background subtraction (RBBS) algorithm implemented in Fiji-ImageJ. CBS often subtracted too much from the fluorescence intensity of G25 aggregates (Figure 2c). In contrast, RBBS maintained the fluorescence intensity of G25 aggregates (Figure 2c). Detection efficiency was evaluated using RBBS, followed by digital intensity amplification. The efficiency of automatic detection of aggregate-positive cells was dramatically improved by combining RBBS with digital intensity amplification (Figure 2d, lanes 1–3). However, as the ball size for RBBS was reduced, the detection efficiency varied from image to image (Figure 2d, lane 4), suggesting that an optimal ball size is required to achieve reproducible detection efficiency. The change in the digital intensity amplification value did not show a dramatic difference compared to that of the pixel size in RBBS. However, since the signal-to-noise ratio is generally important for intensity-based detection of objects, verification remains an inevitable variable in the automatic detection of puncta. Furthermore, the proportion of FPs did not increase when combining RBBS with digital intensity amplification, compared to using no RBBS (Figure 2d, lanes 1–4). However, since a detection efficiency of approximately 100 % was not achieved, further improvements were examined.

**Figure 2.**
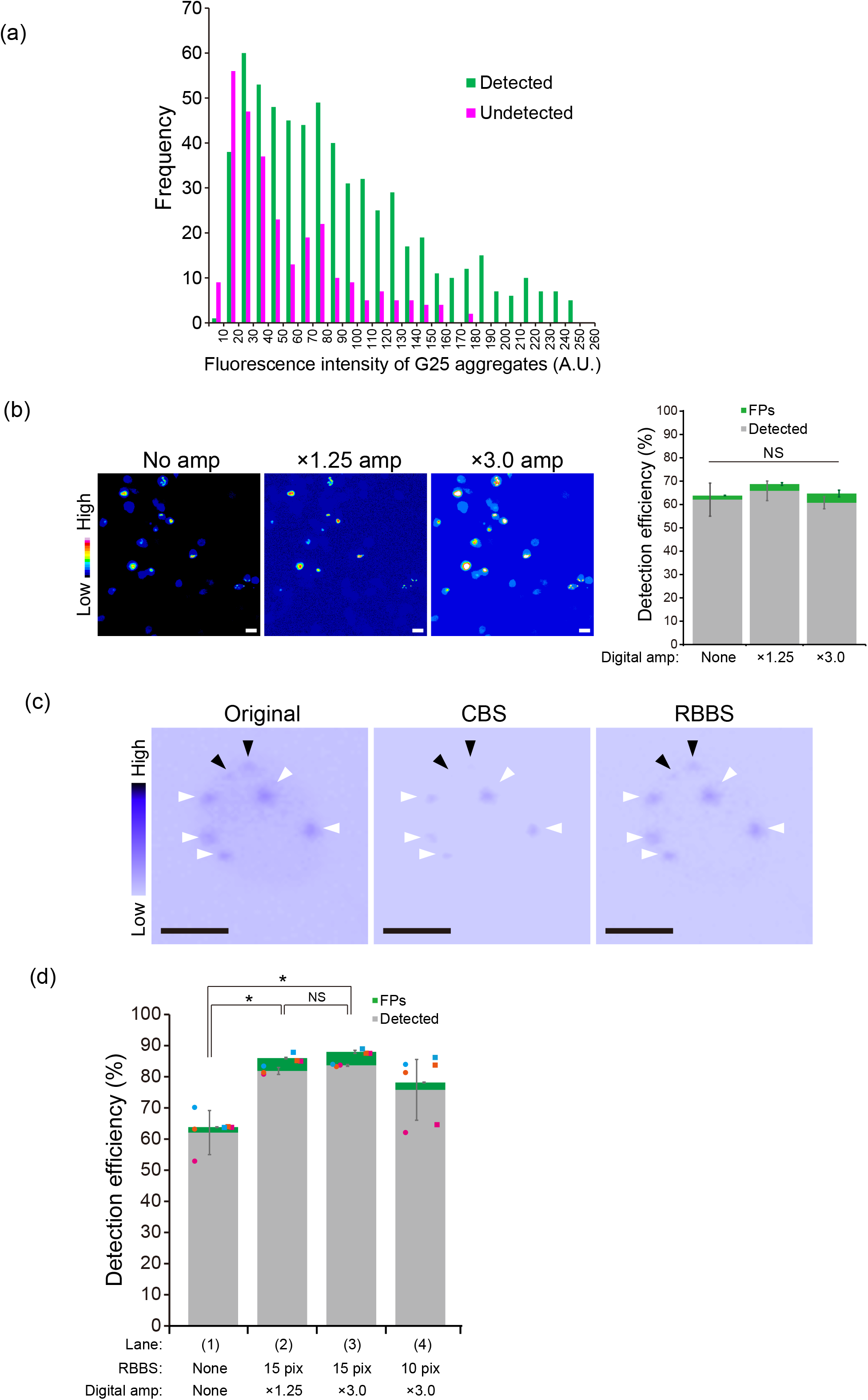
Improving the automatic detection of TDP25 aggregates in Neuro2a cells by the increase in signal-to-noise ratio. (a) Histogram of mean fluorescence intensities of GFP-TDP25 aggregates detected and undetected using PunctaFinder (green and magenta, respectively). (b) (left) Typical LSM images of Neuro2a cells harboring aggregates, with fluorescence intensity digitally amplified (Digital amp: No, ×1.25, and ×3.0 amp.). The color scale on the left of the microscope images represents the pseudocolor look-up table. Bar = 10 μm. (right) Detection efficiency of the detected aggregates (gray) and FPs (green) when the fluorescence intensity was digitally amplified. NS: No significant difference between the three samples (*p* ≥ 0.05). (c) Typical LSM images of a Neuro2a cell that harbors aggregates, where the fluorescence intensity of some aggregates is lost with CBS, but not with RBBS. Original refers to the image without background subtraction. The color scale on the left of the microscope images represents the pseudocolor look-up table. The black and white arrowheads indicate the positions of lost and remaining aggregates, respectively. Bars = 10 μm. (d) Detection efficiency of cells harboring aggregates in the cytoplasm when changing the ball size in RBBS (15 or 10 pixels) and the digital intensity amplification value (×1.25 or ×3.0). The bars and dots show the results from three independent trials (mean ± SD; n = 3). The color of the dots corresponds to the trials. Statistical significance: \**p* < 0.05 and NS: No significant (*p* ≥ 0.05).

### 3.3. Further improvement in the detection efficiency of aggregate-positive cells by reducing the number of target aggregates per analyzed image

We compared the number of aggregate-positive cells detected automatically and manually across eight independently acquired images of G25-expressing cells. The detection efficiency did not change for images containing a high number of aggregate-positive cells, regardless of whether RBBS or digital intensity amplification processing was applied (processed versus unprocessed) (Figure 3a). However, the detection efficiency improved for images containing fewer G25-expressing cells, particularly when the original images were further cropped (Figure 3b). Next, by combining RBBS and digital intensity amplification, we achieved a detection efficiency of 90% as the cumulative value across all trials for images with fewer aggregate-positive cells, compared to a detection efficiency of 78% in unprocessed images (Figure 3b). This suggest that the detection efficiency of puncta may improve when there are fewer cells in a single image. However, because G25 aggregates exhibit a wide range of fluorescence intensities and shapes, images with undetected cells under these conditions tend to lower the overall detection efficiency. Achieving 100% detection efficiency still requires manual correction for undetected cells. However, reducing the number of cells in a single image may simplify the correction process after the automatic detection of aggregate-positive cells using PunctaFinder. Furthermore, when measuring G25 aggregates in nematode embryos, detection efficiencies exceeding 90% without FPs were achieved by combining RBBS and digital intensity amplification, along with reducing the number of embryos analyzed (Figure 3c). Since these embryos originated from parents of a single clone expressing G25, the fluorescence intensity of G25 aggregates within the embryos was relatively consistent, compared to that in cells transiently expressing G25. This consistency highlights that the analytical conditions established here significantly contributed to the detection efficiency of G25 aggregates in both cells and nematode embryos.

**Figure 3.**
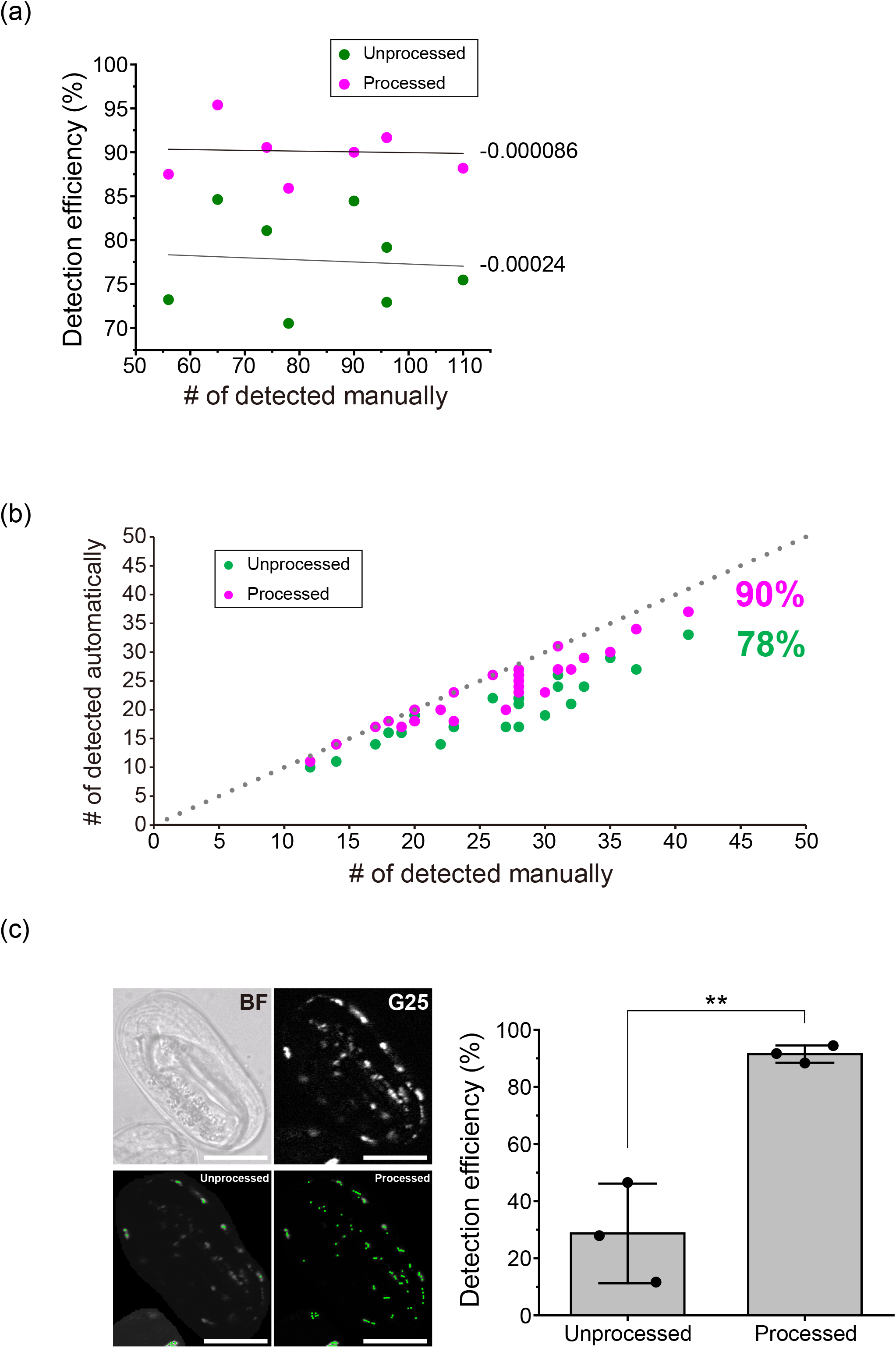
Improving detection accuracy by decreasing the number of detection targets in the image. (a) Correlation between the detection efficiency of Neuro2a cells harboring aggregates of GFP-TDP25 in the cytoplasm and the number of aggregates in a single image. The detection efficiency is shown before and after RBBS and digital intensity amplification processing (Unprocessed and Processed, respectively). The line shows a linear regression, with the regression slopes shown on the right side of the lines. (b) The correlation between the number of Neuro2a cells positive for aggregates as detected automatically and manually, when the number of cells positive for aggregates in an analytical field of view is reduced by cropping the image. The dotted line shows the 100% accuracy (*y* = *x*). The inset values show the total detection efficiency of all processed (green) or unprocessed (magenta) samples. (c) Automatic detection of GFP-TDP25 aggregates in live nematode embryos. Left: Confocal fluorescence and bright-field images of a nematode embryo (top). The green overlayed spots indicate the position of the PunctaFinder-detected aggregates in the embryo in unprocessed and processed confocal fluorescence images (bottom). Bars = 20 μm. Right: Comparison of the detection efficiency of aggregates in nematode embryos between unprocessed and processed confocal fluorescence images. The bars and dots show three independent trials (mean ± SD; n = 3). Statistical significance: \*\**p* < 0.01.

### 3.4. Issues and perspectives for more efficient detection

In this study, we investigated the conditions for effective automated detection of G25 aggregates, which vary in number, size, and fluorescence brightness from cell to cell, using the PunctaFinder algorithm. It is important to note that we achieved a significant improvement in detection accuracy by employing basic image analysis tools, including background removal and brightness amplification, to improve the signal-to-noise ratio. In the original PunctaFinder report, autophagy-related ATG9-positive cells, lipid droplets, and destabilized luciferase aggregates were measured with high precision [14]. This high detection accuracy, achieved without background removal and brightness amplification, can be attributed to the use of images with a small number of cells and relatively uniform fluorescence brightness, due to stable expression in *Saccharomyces cerevisiae* cells. In our previous studies, stable strains expressing G25 did not form aggregates, and G25 aggregates were transiently expressed [15,16]. Image analysis would be simplified if a strain stably expressing G25, capable of forming cytoplasmic aggregates, could be developed.

Other strategies to improve detection efficiency, such as increasing the number of validated puncta cases in the dataset used by the PunctaFinder algorithm, did not lead to significant improvements. Our validation also showed that, although we did not achieve 100% of visually confirmed G25 aggregates, the automatic detection efficiency is high when the purpose is to automatically measure the number of cells with G25 aggregates. As described above, various biological and analytical challenges remain to be addressed for the automated measurement of intracellular aggregates. This study serves as a successful example of detecting G25 aggregates. However, we believe that further studies will contribute not only to advancements in image processing but also to a deeper understanding of the molecular basis of neurodegenerative diseases and the development of therapeutic approaches.

Since spatial image correlation spectroscopy (SICS) is feasible in measuring the average size of aggregates [19,20], combining SICS with the automatic detection algorithm could improve the detection efficiency of aggregates with heterogeneous sizes and numbers. However, these possibilities should be verified in future studies. To detect aggregates with extremely low expression levels and low fluorescence brightness, applying denoising before amplifying the brightness or using the RBBS background removal method is expected to be an effective improvement [21].

## 4. Conclusions

Automated detection of intracellular aggregates with varying sizes, numbers, and brightness in fluorescence images, such as those formed by the transient expression of ALS-associated TDP25, remains challenging. In this study, we demonstrated that appropriate background removal and fluorescence intensity amplification, along with adaptations of the PunctaFinder algorithm, can significantly enhance detection accuracy. In particular, background removal and reducing the number of targets in the analyzed images are important for improving detection accuracy. Currently, fully automating the detection of TDP25 aggregates remains challenging and requires manual correction. In this context, tools like PunctaFinder, which highlight the detected puncta within an image, can be highly useful for assisting with manual corrections during aggregate analysis.

## Supplementary Materials

None.

## Author Contributions

Conceptualization, A.K.; Methodology, S.S., K.S., and A.K.; Software, S.S. and A.K.; Validation, S.S., and A.K.; Formal Analysis, S.S., and A.K.; Investigation, S.S., and A.K.; Resources, K.S. and A.K.; Data Curation, S.S., and A.K.; Writing – Original Draft Preparation, A.K.; Writing – Review & Editing, S.S., and A.K.; Visualization, S.S., and A.K.; Supervision, A.K.; Project Administration, A.K.; Funding Acquisition, A.K.

## Data Availability Statement

The original data presented in the study are openly available in Open Science Framework at DOI: 10.17605/OSF.IO/45ACD (https://osf.io/45acd/; accessed April 3, 2025).

## Acknowledgments

We would like to thank Editage (http://www.editage.jp; accessed April 1, 2025) for English language editing.

---

**Table A1.**

| $w$ | $T_{\text{ratio local}}$ | $T_{\text{ratio global}}$ | $T_{\text{CV}}$ |
| --- | --- | --- | --- |
| 0 | 2.38 | 9.99 | 0.02 |
| 0.1 | 1.00 | 2.79 | 0.35 |
| 0.3 | 1.15 | 1.56 | 0.17 |
| 0.5 | 1.00 | 2.22 | 0.32 |
| 0.65 | 1.06 | 2.22 | 0.32 |
| 1 | 1.06 | 1.80 | 0.32 |
| 2 | 1.03 | 1.80 | 0.32 |

## References

1. Chiti, F.; Dobson, C.M. Protein misfolding, functional amyloid, and human disease. Annual review of biochemistry 2006, 75, 333–366, doi:10.1146/annurev.biochem.75.101304.123901.

2. Takahashi, T.; Kikuchi, S.; Katada, S.; Nagai, Y.; Nishizawa, M.; Onodera, O. Soluble polyglutamine oligomers formed prior to inclusion body formation are cytotoxic. Human molecular genetics 2008, 17, 345–356, doi:10.1093/hmg/ddm311.

3. Kitamura, A.; Nagata, K.; Kinjo, M. Conformational analysis of misfolded protein aggregation by FRET and live-cell imaging techniques. International journal of molecular sciences 2015, 16, 6076–6092, doi:10.3390/ijms16036076.

4. Kim, J.; Lee, H.; Lee, J.H.; Kwon, D.Y.; Genovesio, A.; Fenistein, D.; Ogier, A.; Brondani, V.; Grailhe, R. Dimerization, oligomerization, and aggregation of human amyotrophic lateral sclerosis copper/zinc superoxide dismutase 1 protein mutant forms in live cells. The Journal of biological chemistry 2014, 289, 15094–15103, doi:10.1074/jbc.M113.542613.

5. Shin, Y.; Brangwynne, C.P. Liquid phase condensation in cell physiology and disease. Science 2017, 357, doi:10.1126/science.aaf4382.

6. Jawerth, L.; Fischer-Friedrich, E.; Saha, S.; Wang, J.; Franzmann, T.; Zhang, X.; Sachweh, J.; Ruer, M.; Ijavi, M.; Saha, S.; et al. Protein condensates as aging Maxwell fluids. Science 2020, 370, 1317–1323, doi:10.1126/science.aaw4951.

7. Hartl, F.U. Protein Misfolding Diseases. Annual review of biochemistry 2017, 86, 21–26, doi:10.1146/annurev-biochem-061516-044518.

8. Jaqaman, K.; Loerke, D.; Mettlen, M.; Kuwata, H.; Grinstein, S.; Schmid, S.L.; Danuser, G. Robust single-particle tracking in live-cell time-lapse sequences. Nature methods 2008, 5, 695–702, doi:10.1038/nmeth.1237.

9. Tinevez, J.Y.; Perry, N.; Schindelin, J.; Hoopes, G.M.; Reynolds, G.D.; Laplantine, E.; Bednarek, S.Y.; Shorte, S.L.; Eliceiri, K.W. TrackMate: An open and extensible platform for single-particle tracking. Methods 2017, 115, 80–90, doi:10.1016/j.ymeth.2016.09.016.

10. Shah, S.I.; Ong, H.L.; Demuro, A.; Ullah, G. PunctaSpecks: A tool for automated detection, tracking, and analysis of multiple types of fluorescently labeled biomolecules. Cell calcium 2020, 89, 102224, doi:10.1016/j.ceca.2020.102224.

11. Klickstein, J.A.; Mukkavalli, S.; Raman, M. AggreCount: an unbiased image analysis tool for identifying and quantifying cellular aggregates in a spatially defined manner. The Journal of biological chemistry 2020, 295, 17672–17683, doi:10.1074/jbc.RA120.015398.

12. Ershov, D.; Phan, M.S.; Pylvanainen, J.W.; Rigaud, S.U.; Le Blanc, L.; Charles-Orszag, A.; Conway, J.R.W.; Laine, R.F.; Roy, N.H.; Bonazzi, D.; et al. TrackMate 7: integrating state-of-the-art segmentation algorithms into tracking pipelines. Nature methods 2022, 19, 829–832, doi:10.1038/s41592-022-01507-1.

13. Mantes, A.D.; Herrera, A.; Khven, I.; Schlaeppi, A.; Kyriacou, E.; Tsissios, G.; Skoufa, E.; Santangeli, L.; Buglakova, E.; Durmus, E.B.; et al. Spotiflow: accurate and efficient spot detection for fluorescence microscopy with deep stereographic flow regression. bioRxiv 2024, 2024.2002.2001.578426, doi:10.1101/2024.02.01.578426.

14. Terpstra, H.M.; Gomez-Sanchez, R.; Veldsink, A.C.; Otto, T.A.; Veenhoff, L.M.; Heinemann, M. PunctaFinder: An algorithm for automated spot detection in fluorescence microscopy images. Molecular biology of the cell 2024, 35, mr9, doi:10.1091/mbc.E24-06-0254.

15. Kitamura, A.; Nakayama, Y.; Shibasaki, A.; Taki, A.; Yuno, S.; Takeda, K.; Yahara, M.; Tanabe, N.; Kinjo, M. Interaction of RNA with a C-terminal fragment of the amyotrophic lateral sclerosis-associated TDP43 reduces cytotoxicity. Scientific reports 2016, 6, 19230, doi:10.1038/srep19230.

16. Fujimoto, A.; Kinjo, M.; Kitamura, A. Short Repeat Ribonucleic Acid Reduces Cytotoxicity by Preventing the Aggregation of TDP-43 and Its 25 KDa Carboxy-Terminal Fragment. JACS Au 2024, 4, 3896–3909, doi:10.1021/jacsau.4c00566.

17. Kitamura, A.; Fujimoto, A.; Kawashima, R.; Lyu, Y.; Sasaki, K.; Hamada, Y.; Moriya, K.; Kurata, A.; Takahashi, K.; Brielmann, R.; et al. Hetero-oligomerization of TDP-43 carboxy-terminal fragments with cellular proteins contributes to proteotoxicity. Commun Biol 2024, 7, 743, |p|doi:10.1038/s42003-024-06410-3.

18. Brenner, S. The genetics of behaviour. Br Med Bull 1973, 29, 269–271, doi:10.1093/oxfordjournals.bmb.a071019.

19. Hebert, B.; Costantino, S.; Wiseman, P.W. Spatiotemporal image correlation spectroscopy (STICS) theory, verification, and application to protein velocity mapping in living CHO cells. Biophysical journal 2005, 88, 3601–3614, doi:10.1529/biophysj.104.054874.

20. Kitamura, A.; Shimizu, H.; Kinjo, M. Determination of cytoplasmic optineurin foci sizes using image correlation spectroscopy. Journal of biochemistry 2018, 164, 223–229, doi:10.1093/jb/mvy044.

21. Li, X.; Li, Y.; Zhou, Y.; Wu, J.; Zhao, Z.; Fan, J.; Deng, F.; Wu, Z.; Xiao, G.; He, J.; et al. Real-time denoising enables high-sensitivity fluorescence time-lapse imaging beyond the shot-noise limit. Nature biotechnology 2023, 41, 282–292, doi:10.1038/s41587-022-01450-8.

